# Engineered hydrogel reveals contribution of matrix mechanics to esophageal adenocarcinoma 3D organoids and identify matrix-activated therapeutic targets

**DOI:** 10.1101/2022.11.13.516357

**Authors:** Ricardo Cruz-Acuña, Secunda W. Kariuki, Kensuke Sugiura, Claudia Loebel, Tatiana Karakasheva, Joel T. Gabre, Jason A. Burdick, Anil K. Rustgi

## Abstract

Increased extracellular matrix (ECM) stiffness has been implicated in esophageal adenocarcinoma (EAC) progression, metastasis, and resistance to therapy. However, the underlying pro-tumorigenic pathways are yet to be defined. Additional work is needed to develop physiologically relevant *in vitro* 3D culture models that better recapitulate the human tumor microenvironment and can be used to dissect the contributions of matrix stiffness to EAC pathogenesis. Here, we describe a modular, tumor ECM-mimetic hydrogel platform with tunable mechanical properties, defined presentation of cell-adhesive ligands, and protease-dependent degradation that supports robust *in vitro* growth and expansion of patient-derived EAC 3D organoids (EAC PDOs). Hydrogel mechanical properties control EAC PDO formation, growth, proliferation and activation of tumor-associated pathways that elicit stem-like properties in the cancer cells, as highlighted through *in vitro* and *in vivo* environments. We also demonstrate that the engineered hydrogel serves as a platform to identify potential therapeutic targets to disrupt the contribution of pro-tumorigenic increased matrix mechanics in EAC. Together, these studies show that an engineered PDO culture platform can be used to inform the development of therapeutics that target ECM stiffness in EAC.

## Introduction

Over the past 30 years the incidence of esophageal adenocarcinoma (EAC) has risen dramatically by 300-600% in the US^1^. The pathogenesis of EAC involves epigenetic, genomic, and genetic alterations and an interplay with microenvironmental changes^2–5^. Previous studies have demonstrated that changes in the tumor microenvironment, involving a stiffened extracellular matrix (ECM), are associated with EAC progression^6–10^. Although clinical observations suggest that increased ECM stiffness drive cell transformation, cancer progression and metastasis, the underlying pathways of mechanotransduction that lead cancer cells to translate mechanical signals into intracellular pro-tumorigenic pathways are yet to be defined^9^. Therefore, additional work is needed to develop physiologically relevant 3D culture models that better recapitulate the human tumor microenvironment and can dissect the contributions of matrix properties to elucidate underlying molecular mechanisms of the disease^11^.

Patient-derived tumor organoids have become attractive pre-clinical models to study cancer biology as they retain the biological characteristics of the primary tumor^11–13^. Indeed, our lab has shown that patient-derived esophageal adenocarcinoma organoids (EAC PDOs) can serve as avatars to study cellular responses to anticancer drugs, as they recapitulate patient’s drug responses in the clinic^14,15^. Patient-derived organoids (PDOs) are traditionally grown in Matrigel™, a heterogeneous, complex mixture of ECM proteins, proteoglycans and growth factors secreted by Engelbreth-Holm-Swarm mouse sarcoma cells^16^. However, Matrigel™ suffers from lot-to-lot compositional and structural variability, and cannot recapitulate the independent role of matrix properties to disease progression due to the inability to uncouple its physicochemical properties^16–18^. For instance, changes to the bulk concentration (e.g. decrease in matrix density) of Matrigel™ is a common approach to vary its mechanical properties, however, these changes unavoidably alter other matrix properties, such as adhesive ligand density and fiber density/structure^18^ (Supplementary Fig. 1a). Therefore, although modulation of the bulk concentration of Matrigel™ results in changes in EAC PDO formation (density), growth (area), and transcriptional expression of EAC-associated genes (Supplementary Fig. 1b-d), it is unclear whether this effect is mediated by differences in mechanical or biochemical matrix properties. To address this important gap, well-defined engineered hydrogels are an evolving and important component of tumor PDO culture systems as alternatives to Matrigel™, particularly to introduce user-defined microenvironment signals to study human epithelial tumors^19–23^.

Here, we describe a modular, tumor ECM-mimetic hydrogel platform with defined physicochemical properties that support EAC PDO culture and growth. Hydrogel mechanical properties, adhesive ligand presentation, and protease-dependent degradation were key parameters in engineering a hydrogel that supported EAC PDO viability and growth. Particularly, hydrogel mechanical properties controlled EAC PDO formation and growth, and activation of tumor-associated pathways that elicit stem-like properties in the cancer cells as highlighted through *in vitro* and *in vivo* models. Additionally, the engineered hydrogel served as a platform to identify potential therapeutic targets to disrupt the contribution of pro-tumorigenic increased matrix mechanics in EAC. Whereas previous work has established engineered hydrogels as tumor ECMs to investigate multicellular assembly and tumor invasion using cancer cell lines^24–26^, or study tumor PDO resistance to therapy^23,27–29^, we are the first to analyze the novel contributions of ECM mechanical properties to EAC PDO growth, proliferation, and identification of matrix mechanics-mediated drivers of stem-like properties as therapeutic targets. Finally, the modular nature of the engineered hydrogel platform allows for potential adaption to the culture of 3D organoid models of other human cancers. Thus, provide new mechanistic and translational insights with broad applicability.

## Results

### Engineered hydrogel supports EAC PDO development

We selected a hydrogel platform based on hyaluronic acid (HA), specifically through the crosslinking of norbornene HA (NorHA) macromer (Supplementary Fig. 2a), which exhibits native bio-functionality and has been extensively developed for *in vitro* cell activation by stiffening events and several pre-clinical *in vivo* applications^30^. HA has inherent biological importance due to its binding to cell receptors (e.g. CD44^31,32^) and is a major component of the tumor niche (Fig. 1a,b), creating a microenvironment that is favorable for tumor angiogenesis, invasion, and metastasis^33,34^. Certainly, EAC patient biopsies showed increased expression of HA in the tumor microenvironment (Fig. 1a), whereas RNA-sequencing analysis of 286 esophageal cancer and 283 normal esophageal tissues collected from The Cancer Genome Atlas (TCGA) and The Genome-Tissue Expression Project (GTEx)^35^ confirmed increased expression of HA synthesis genes (HAS1, HAS2, UDGH) in the tumor microenvironment (Fig. 1b). Moreover, our hydrogel system offers significant advantages due to its well-defined structure, covalent incorporation of peptide sequences for enhanced cell–matrix interactions, and user-defined hydrogel stiffness by varying the crosslinking peptide concentration, which mediates crosslinking via a thiol-norbornene reaction to form a NorHA hydrogel (Fig. 1c and Supplementary Fig. 2a)^36^. Indeed, HA-based hydrogels have been used by other groups to study tumor progression and resistance to therapy of other cancer types (e.g. colorectal and pancreatic adenocarcinomas)^28,29,37^.

**Figure 1:**
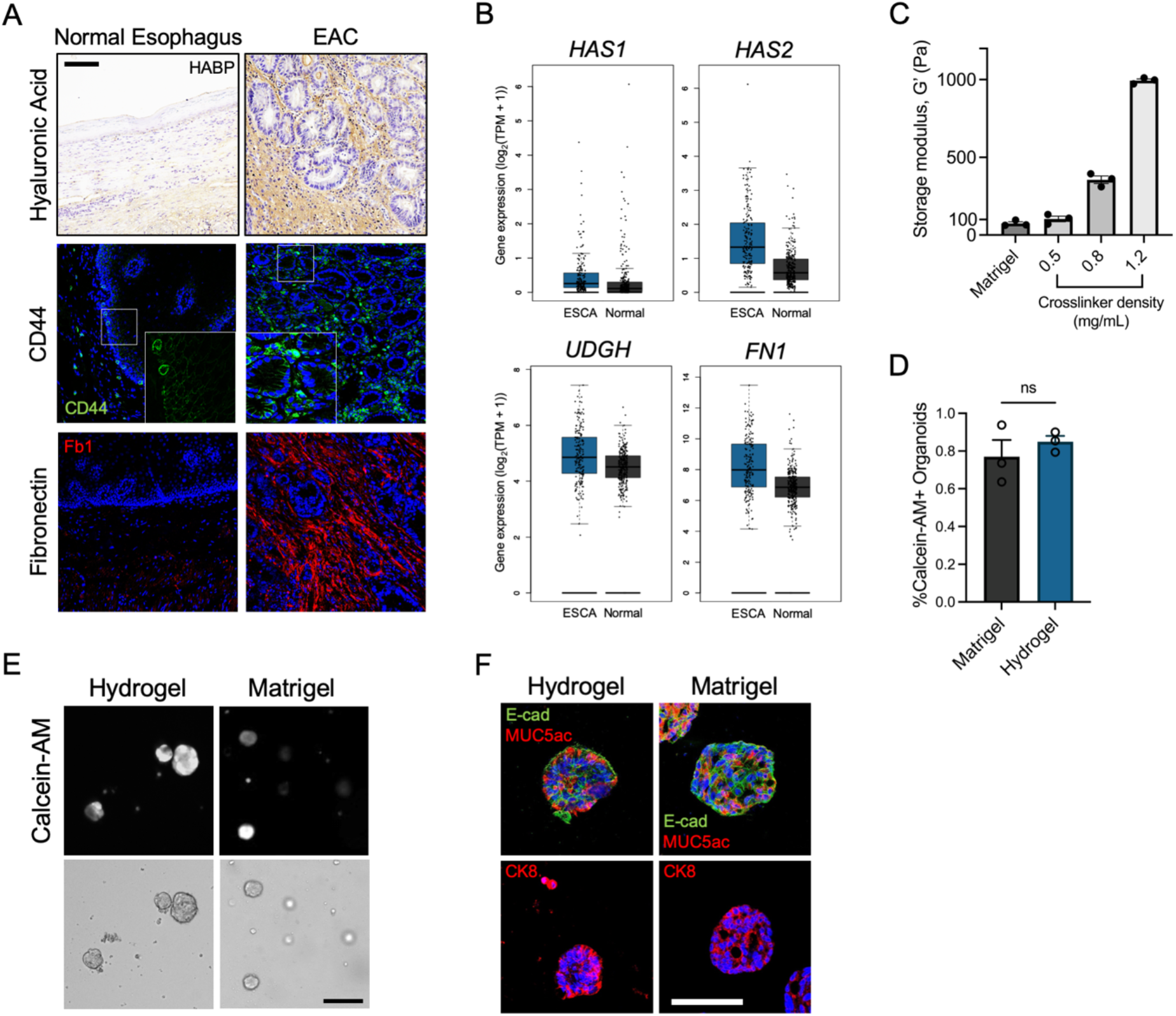
Engineered hydrogel supports EAC PDO development. (A) Images of patient tissue sections from normal or EAC biopsies stained for hyaluronic acid (HABP), CD44, or Fibronectin (Fb1). Scale bar, 100 µm. (B) Bulk RNA sequencing analysis of esophageal carcinoma (ESCA) and normal pancreatic tissue samples for fibronectin (FN1) and hyaluronan associated genes (HAS1, HAS2, UGDH). n = 286 for ESCA, and n = 283 for Normal. (C) Relationship between crosslinker density (mg/mL) and storage modulus, G’ (mean ± SEM; n = 3 independently prepared hydrogels per condition). (D) Quantification of PDO viability as assessed by Calcein-AM labelling at 7 days after encapsulation. Viability is quantified as the percentage of PDOs that stained positive for Calcein-AM (mean ± SEM; n = average number of Calcein-AM+ organoids per hydrogel; at least 20 organoids per hydrogel were analyzed). Welch’s t-test with two-tailed comparison showed no significant differences between groups (ns = P > 0.05). (E) Representative transmitted light and fluorescence microscopy images of EAC PDOs cultured in NorHA hydrogels or Matrigel. Scale bars, 200 µm. (F) Representative fluorescence microcopy images of EAC PDOs within NorHA hydrogels stained for Mucin 5AC (MUC5ac), e-cadherin (e-cad), and Cytokeratin 8 (CK8) at 14 days post-encapsulation. Scale bar: 50 µm. Three independent experiments were performed and data are presented for one of the experiments. Every independent experiment was performed with four gel samples per experimental group.

In our study, we explored a NorHA hydrogel formulation that supports the viability of EAC PDOs generated in Matrigel™. After EAC PDOs were grown in Matrigel™, they were retrieved, dissociated into single cells, and encapsulated in NorHA hydrogels with similar mechanical properties as Matrigel™ (G’ = 100 Pa; Fig. 1c and Supplementary Fig. 2b). NorHA hydrogels were engineered to present a constant 2.0 mM RGD adhesive peptide (GCGYGRGDSPG) density and crosslinked with the protease-degradable peptide VPM (0.5 mg/mL; GCNSVPMSMRGGSNCG). Incorporation of 2.0 mM RGD adhesive peptide and a protease-degradable crosslinker to engineered hydrogels has previously promoted epithelial organoid development^17,38,39^. Moreover, the RGD adhesive ligand (α5β1 and ανβ3 integrin binding peptide) is found in many adhesive proteins, including fibronectin that is a major ECM protein component in EAC (Fig. 1a, b)^6,40^. Indeed, EAC patient biopsies showed increased expression of fibronectin (Fig. 1a), and RNA-sequencing analysis of tissues collected from TCGA and GTEx^35^ confirmed increased expression of fibronectin (FN1) in the tumor microenvironment (Fig. 1b). EAC PDOs grown in NorHA hydrogels functionalized with RGD and crosslinked with VPM demonstrated viability comparable to those grown in Matrigel™ (Fig. 1d,e). However, when EAC PDOs were grown in NorHA hydrogels presenting an inactive scrambled peptide (RDG), or functionalized with RGD and crosslinked with non-degradable agent DTT (1,4-dithiothreitol), EAC PDOs showed reduced viability at seven days post-encapsulation, as compared to hydrogels functionalized with RGD and crosslinked with VPM (Supplementary Fig. 3a-c). Moreover, PDOs cultured in the engineered hydrogel formulation showed expression of the epithelial marker e-cadherin (e-cad), and of EAC-specific markers, Mucin 5AC (MUC5ac) and Cytokeratin 8 (CK8) to similar levels as PDOs cultured in Matrigel™ (Fig. 1f). Finally, to test whether NorHA hydrogels are suitable for the culture of other human PDOs, we embedded Barrett’s esophagus (a precursor or premalignant condition that predisposes to EAC) PDOs (BE PDOs)^41^ in the engineered NorHA hydrogel (Supplementary Fig. 3d). BE PDOs cultured in the engineered matrix maintain high viability and comparable growth to the same PDOs cultured in Matrigel™ twelve days after encapsulation (Supplementary Fig. 3d). Taken together, these data suggest the requirement of specific matrix properties that are essential for organoid viability and formation, establishing the engineered NorHA hydrogel as a novel culture system for EAC PDOs that has the potential to be adapted for the generation of different human tumor organoids.

### Matrix mechanics control EAC PDO development

As ECM mechanical properties influence epithelial cell behavior^38,42^, we investigated the influence of crosslinker density, which controls hydrogel mechanical properties (Fig. 1c), on EAC PDO size, formation, and proliferation (Fig. 2a-d). EAC PDOs were embedded in NorHA hydrogels with mechanical properties that ranged from a ‘soft’ hydrogel (0.5 mg/mL VPM; G’ = 100 Pa, similar to Matrigel™) to a ‘stiff’ hydrogel (1.2 mg/mL; G’ = 1000 Pa, similar to tumor ECM), and cultured for 14 days. Importantly, the mechanical properties of our stiff hydrogel have been shown to promote pro-tumorigenic behavior in normal cells in previous *in vitro* studies^43,44^, and compare well with measurements of human tumor stiffness^43,45,46^. EAC PDOs embedded in stiff (G’: 1,000 Pa) NorHA hydrogels showed significant increases in the size (area) and formation (density) of organoids formed per hydrogel as a function of matrix stiffness (Fig. 2a-c). Similarly, PDOs embedded in the stiff hydrogel condition showed increased cell proliferation, as compared to organoids embedded in the soft hydrogel condition (Fig. 2c,d). These data suggest that a stiff matrix nurtures EAC PDO growth, formation, and proliferation as compared to a soft matrix, establishing the engineered hydrogel as an innovative and exciting platform to investigate the independent contributions of matrix mechanics in EAC PDO development.

**Figure 2:**
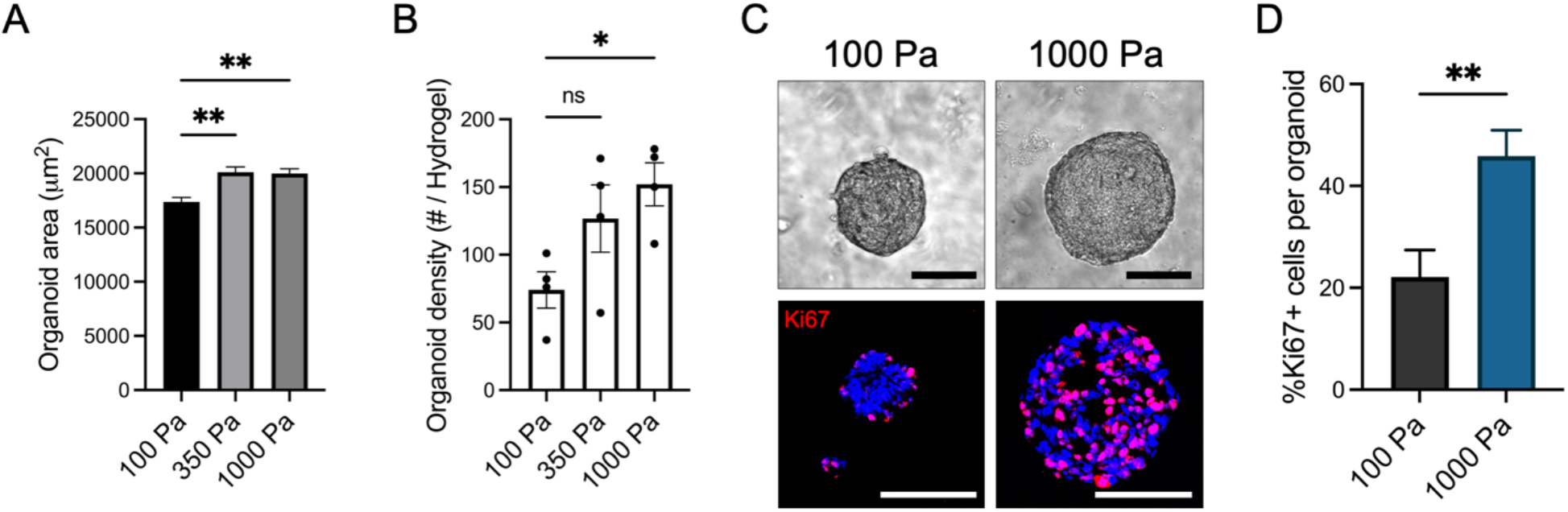
Engineered hydrogel stiffness modulates EAC PDO development. (A) Quantification of PDO (A) size (area) and (B) density as a function of matrix stiffness at 14 days post-encapsulation (mean ± SEM; (A) n = at least 300 organoids analyzed across 4 hydrogels per group; (B) n = 4 hydrogels per group). (A, B) Kruskal-Wallis test with Dunn’s multiple comparisons showed significant differences between 100 Pa and 350 Pa or 1000 Pa (*P <0.05, **P <0.01). (C) Representative transmitted light and fluorescence microscopy images of EAC PDOs, and (D) quantification of proliferating cells (%Ki67+) in EAC PDOs cultured in NorHA hydrogels of different stiffnesses at 14 days post-encapsulation (mean ± SEM; n = at least 15 organoids analyzed per group). Mann-Whitney test showed significant differences between 100 Pa and 1000 Pa (**P <0.01). Scale bar: 100 µm. (A-D) Three independent experiments were performed and data are presented for one of the experiments. Every independent experiment was performed with four gel samples per experimental group.

### Matrix mechanics modulates YAP activation in EAC PDOs

Recent work has demonstrated that dysregulated Yes-associated protein 1 (Yap) activation is essential for the growth of most solid tumors by inducing cancer stem cell features, proliferation, and metastasis^47–49^. Dysregulated Yap activation is a major determinant of stem cell properties by direct upregulation of SOX9^50,51^ in EAC. However, the pathophysiological event(s) that elicits upregulation of the YAP-SOX9 axis in EAC remain elusive. Therefore, as Yap functions as a sensor of the structural and mechanical features of the cell microenvironment, we investigated whether changes in matrix biomechanics played a role in the expression of Yap and Sox9 in EAC PDOs. EAC PDOs embedded in the stiff hydrogel showed a significant increase in YAP and Sox9 expression as compared to the same organoids within the soft hydrogel or Matrigel™ (Fig. 3a, b and Supplemental Fig. 4a, b). Similarly, protein expression of the esophageal cancer stem cell marker CD44^52,53^ was significantly higher in EAC PDOs within the stiff hydrogel as compared to organoids within the soft hydrogel or Matrigel™ (Fig. 3c). Moreover, the expression of other EAC-associated genes (namely, P53 and STAT3) known to respond to mechanical cues significantly increased in EAC PDOs within the stiff hydrogel as compared to the same organoids within the soft hydrogel or Matrigel™ (Supplementary Fig 5a). These data suggest that increased matrix stiffness promotes aberrant activation of the YAP-SOX9 axis, endowing stem-like properties to the EAC PDOs that are associated with increased PDO growth, formation, cell proliferation and CD44 expression.

**Figure 3:**
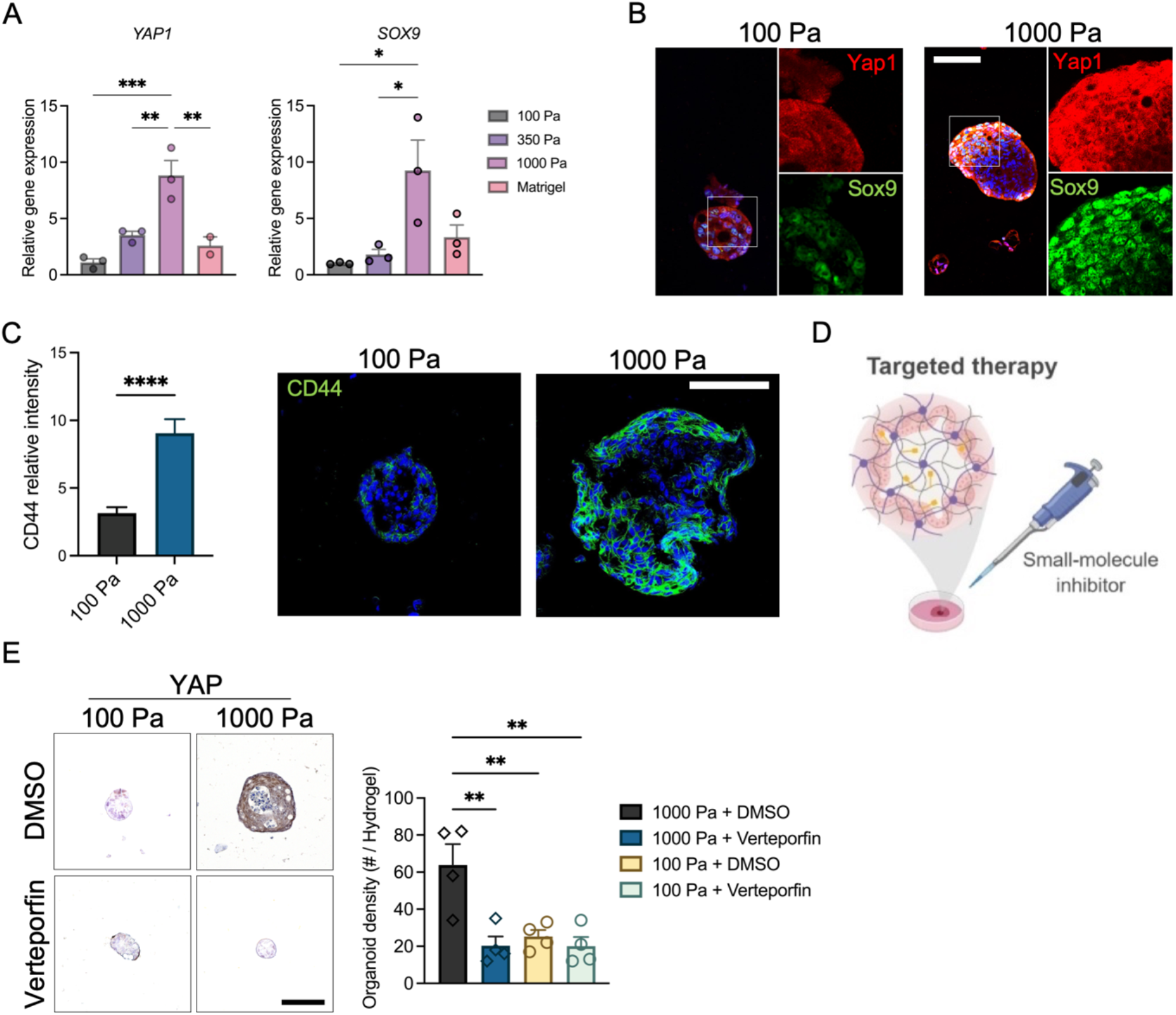
Engineered hydrogel stiffness modulates YAP activation in EAC PDOs. (A) Transcriptional expression and (B) representative fluorescence images of YAP and SOX9 in organoids within NorHA hydrogels of different stiffness at 14 days post-encapsulation (mean ± SEM; n = 3 technical replicates, representative of 3 independent experiments). (A) One-way ANOVA with Tukey’s multiple comparisons test showed significant differences between 100 Pa and 1000 Pa, 350 Pa and 1000 Pa, 1000 Pa and Matrigel™ (*P <0.05, **P <0.01, ***P <0.001). (B) RNA levels normalized to 100 Pa. Scale bar: 100 µm. (C) Quantification and representative fluorescence microscopy images of CD44 expression in EAC PDOs cultured in NorHA hydrogels of different stiffnesses at 14 days post-encapsulation (mean ± SEM; n = at least 20 organoids analyzed per group). Mann-Whitney test showed significant differences between 100 Pa and 1000 Pa (****P <0.0001). (D) Schematic of *in vitro* experiment of EAC PDOs within NorHA hydrogels being treated with YAP inhibitor, Verteporfin. Created with BioRender.com. (E) Representative immunohistochemistry microscopy images and quantification of Yap expression in EAC PDOs cultured in NorHA hydrogels of different stiffnesses at 7 days post-encapsulation and treated with 20 nM Verteporfin or DMSO (mean ± SEM; n = 4 hydrogels per group). One-way ANOVA with Tukey’s multiple comparisons test showed significant differences between 1000 Pa + DMSO and every other group (**P <0.01), and no significant differences among other groups (P > 0.05). Scale bar: 100 µm. (A-E) Three independent experiments were performed and data are presented for one of the experiments. Every independent experiment was performed with four gel samples per experimental group.

Another advantage of HA-based hydrogels is its permissive diffusional properties, which allow diffusion of small molecules, including drugs and inhibitors, to cells (Fig. 3d)^30,54^. Therefore, we investigated if the effect of increased matrix stiffness on EAC PDO was repressed via inhibition of the nuclear translocation of Yap with Verteporfin, a commercially-available small molecule inhibitor whose efficacy and potential as therapy has been described previously^55,56^. Addition of Verteporfin to the cell culture media of EAC PDOs grown in NorHA hydrogels (Fig. 3d) resulted in significant reduction in Yap expression, and the formation (density) and size of organoids within the stiff hydrogel, as compared to the vehicle control (DMSO; Fig. 3e and Supplementary Fig. 4c). However, EAC PDOs embedded in the soft hydrogel and exposed to Verteporfin showed no significant differences in Yap expression, organoid formation and size as compared to vehicle control (Fig. 3e and Supplementary Fig. 4c). These data further underscore that matrix mechanics modulate the dysregulated activation of YAP-SOX9. Finally, these results suggest that NorHA hydrogels can serve as an engineered organoid culture platform to investigate the effect of matrix mechanics in the expression of tumor-associated genes and identification of potential therapeutic targets.

### Matrix mechanics control EAC PDO development and YAP activation *in vivo*

Engineered hydrogels have been utilized previously as organoid delivery vehicles^30,39^. Therefore, we embedded organoids in our engineered tumor ECM-mimetic hydrogels and transplanted into dorsal subcutaneous spaces of immunocompromised mice to study the effects of matrix mechanics (Fig. 4a). After 4 weeks, PDOs showed expression of the epithelial marker e-cadherin (e-cad), and of EAC-specific markers, Cytokeratin 8 (CK8), and Mucin 5AC (MUC5ac; Fig. 4b). However, implanted stiff hydrogels contained significantly larger (area) EAC PDOs as compared to organoids within soft hydrogels (Fig. 4c and Supplementary Fig. 4d). Additionally, PDOs embedded in the stiff hydrogel showed increased cell proliferation, Sox9 expression and nuclear localization of YAP, as compared to organoids embedded within the soft hydrogel (Fig. 4d, e). These data suggest that matrix mechanics control PDO growth and proliferation via dysregulated activation of the YAP-SOX9 axis in an *in vivo* environment. Finally, as studies suggest that increased ECM stiffness stabilizes mutant p53^57^, and we have previously shown that mutant p53-Yap interactions promote esophageal cancer progression^58^, we investigated p53 expression in EAC PDOs within NorHA hydrogels. We observed that EAC PDOs within the stiff hydrogel show increased nuclear p53 expression, as compared to organoids within the soft hydrogel (Supplementary Fig. 5b). These data establish the engineered hydrogel as a platform to study the contributions of matrix mechanics in the activation of regulatory mechanisms in EAC PDOs.

**Figure 4:**
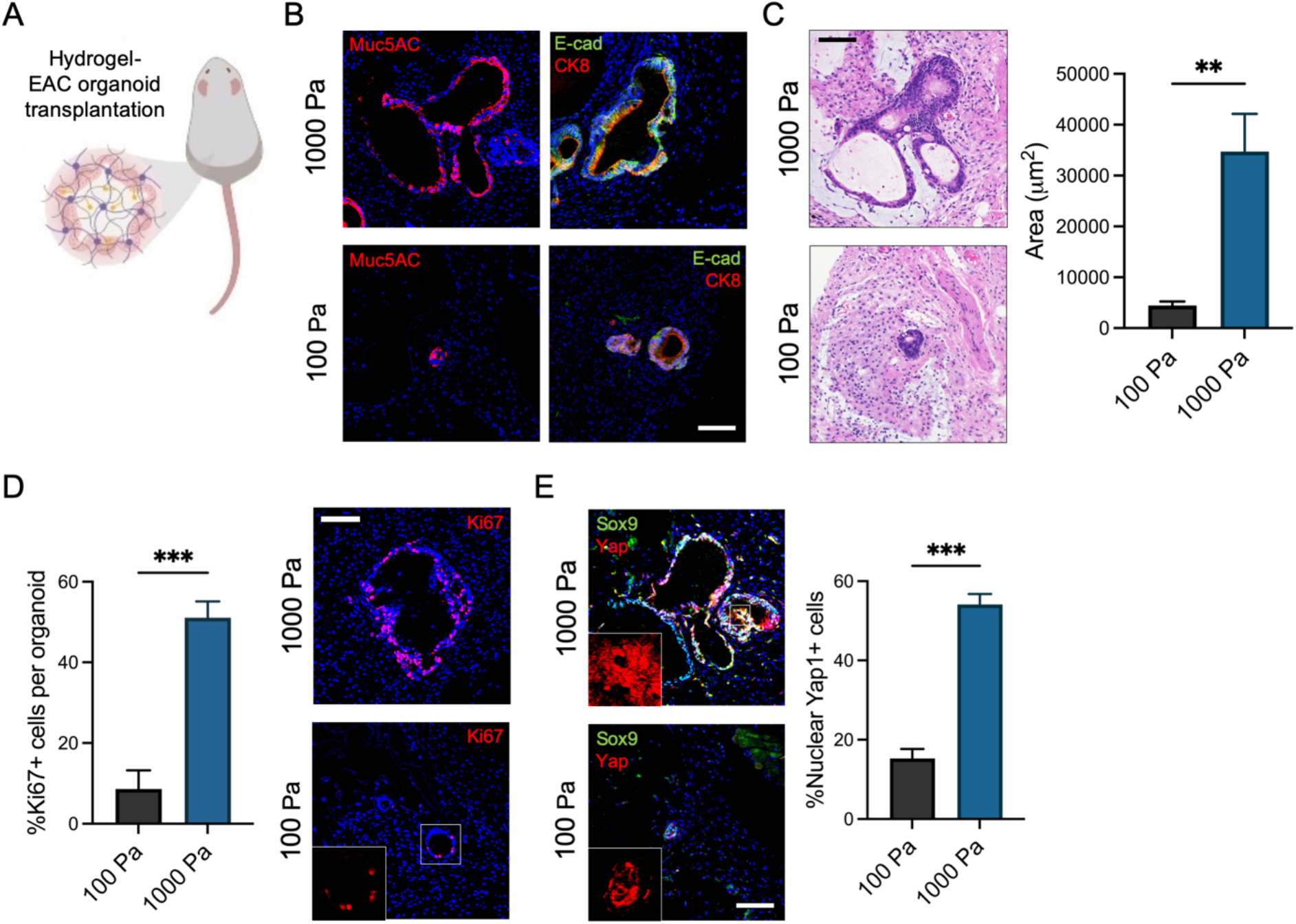
Engineered hydrogel stiffness-dependent growth of EAC PDOs in *in vivo* xenograft model. (A) Schematic of *in vivo* transplantation experiment of EAC PDOs within NorHA hydrogels into mouse subcutaneous pockets. Created with BioRender.com. (B) Representative fluorescence microcopy images of EAC PDOs within NorHA hydrogels stained for Mucin 5AC (MUC5ac), e-cadherin (e-cad), and Cytokeratin 8 (CK8) at 28 days post-encapsulation and *in vivo* transplantation. Scale bar: 100 µm. (C) Histological (H&E) microcopy images and quantification of PDO size (area) as a function of matrix stiffness at 28 days post-encapsulation and *in vivo* transplantation (mean ± SEM; n = at least 5 organoids analyzed across). Welch’s t-test with two-tailed comparison showed significant differences between 100 Pa and 1000 Pa (**P <0.01). Scale bar: 100 µm. (D) Quantification and representative fluorescence microscopy images of percentage of proliferating cells (%Ki67+) per EAC PDOs as a function of matrix stiffness at 28 days post-encapsulation and *in vivo* transplantation (mean ± SEM; n = at least 6 organoids analyzed per group). Mann-Whitney test showed significant differences between 100 Pa and 1000 Pa (***P <0.001). Scale bar: 100 um. (E) Representative fluorescence microcopy images of EAC PDOs within NorHA hydrogels stained for Sox9, and Yap at 28 days post-encapsulation and *in vivo* transplantation. Quantification of percentage of nuclear Yap+ cells (%Yap+) per EAC PDO as a function of matrix stiffness (mean ± SEM; n = 6 organoids analyzed per group). Welch’s t-test with two-tailed comparison showed significant differences between 100 Pa and 1000 Pa (***P <0.001). Scale bar: 100 um. (A-E) Two independent experiments were performed and data are presented for one of the experiments. Every independent experiment was performed with two gels per mouse and five mice per experimental group.

### Yap inhibition represses the effect of matrix mechanics in EAC PDOs *in vivo*

Patient-derived xenograft (PDX) models have proven to be highly effective in predicting the efficacy of both conventional and novel anti-cancer therapeutics^59^. Therefore, we exploited the native biocompatibility of the engineered NorHA hydrogel and the susceptibility of EAC PDOs to anti-cancer drugs^14^, and applied an *in vivo* xenograft model for targeted therapy. EAC PDOs were embedded in soft or stiff NorHA hydrogels and transplanted into the dorsal subcutaneous space of immunocompromised mice. After 1 week, mice were treated with intraperitoneal injections of Verteporfin or vehicle control (DMSO) for 3 weeks (Fig. 5a). At the end of the treatment, EAC PDOs in stiff hydrogels from mice treated with Verteporfin were significantly lower in density and smaller in size as compared to organoids within implanted stiff hydrogels from the control (DMSO) group (Fig. 5b-d and Supplementary Fig. 4e). Additionally, PDOs within the stiff hydrogel from mice treated with Verteporfin showed a significant decrease in Yap and Sox9 expression, and cell proliferation, as compared to organoids within stiff hydrogels from control (DMSO) mice (Fig. 5e-g). Interestingly, EAC PDOs within implanted soft hydrogels from mice treated with Verteporfin showed no significant difference in organoid density and size as compared to organoids within implanted soft hydrogels from control (DMSO) mice (Fig. 5b-d and Supplementary Fig. 4e). Concurrently, when comparing PDOs within the soft hydrogel from mice treated with Verteporfin to those of control (DMSO) mice, organoids showed no significant difference in YAP and Sox9 expression, and cell proliferation (Fig. 5e-g). These data suggest that the effect of increased matrix mechanics in PDO growth, formation, and proliferation is mediated in part by matrix stiffness-dependent activation of YAP-SOX9 in an *in vivo* environment. These data further suggest that matrix stiffness-mediated activation of YAP-SOX9 in EAC PDOs elicit stem-like properties via increased organoid formation, growth and proliferation. Finally, these results suggest that NorHA hydrogels can serve as a platform for the identification of therapeutic targets in patient-derived tumor organoid xenografts studies.

**Figure 5:**
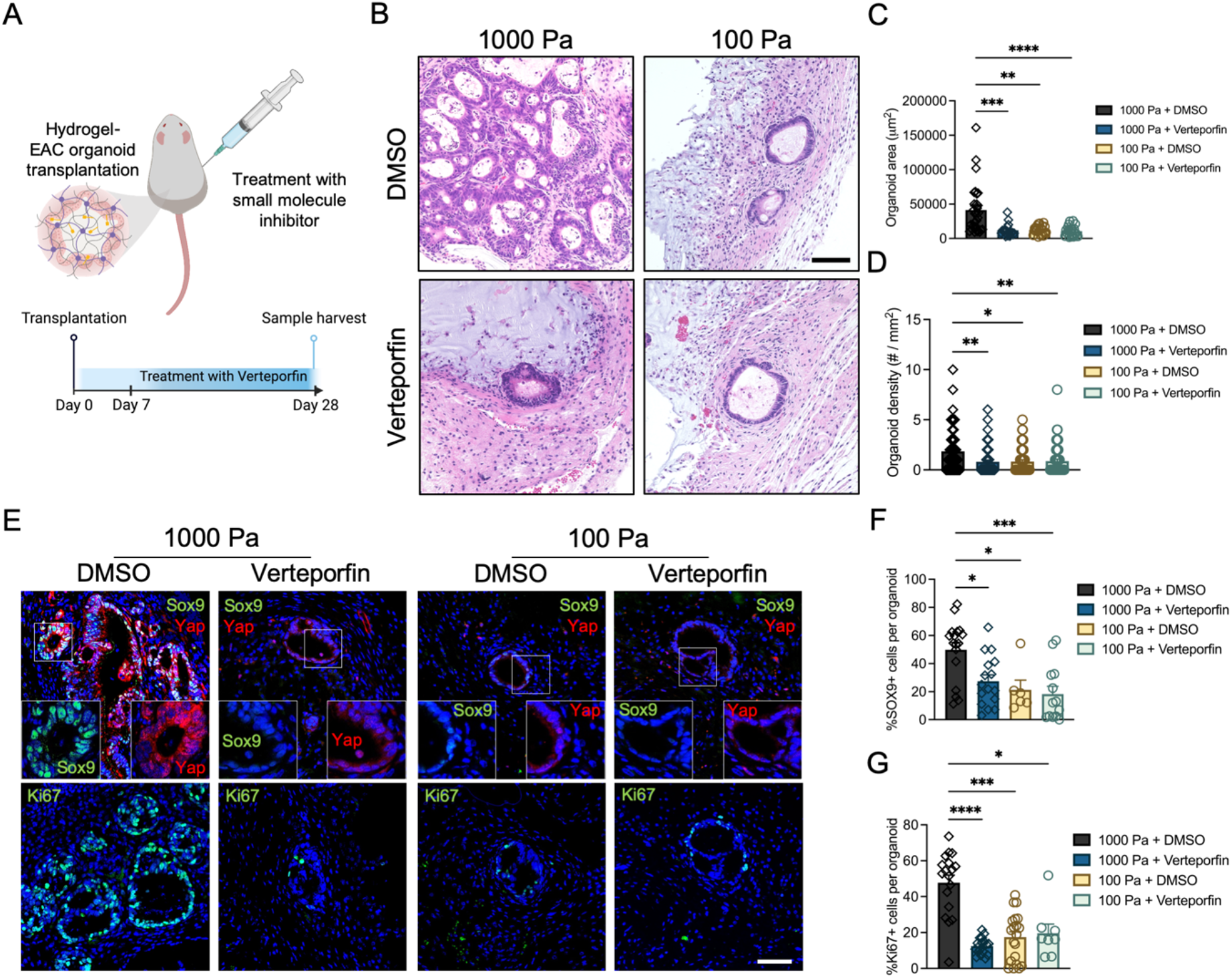
Yap inhibition hinders effect of matrix stiffness in EAC PDOs *in vivo* xenograft model. (A) Schematic of *in vivo* transplantation experiment of EAC PDOs within NorHA hydrogels into mouse subcutaneous pockets and treatment with intraperitoneal injections of Verteporfin or DMSO. Created with BioRender.com. (B) Histological (H&E) microcopy images and quantification of PDO (C) size (area) and (D) density within NorHA hydrogels at 28 days post-encapsulation, *in vivo* transplantation, and treatment with Verteporfin or DMSO (mean ± SEM; (C) n = at least 17 organoids analyzed across 10 samples per group; (D) n = at least 55 organoids analyzed across 10 samples per group). Kruskal-Wallis test with Dunn’s multiple comparisons showed significant differences between 1000 Pa + DMSO and 1000 Pa + Verteporfin, 1000 Pa + DMSO and 100 Pa + DMSO, 1000 Pa + DMSO and 100 Pa + Verteporfin (*P <0.05, **P <0.01, ***P <0.001), and no significant differences among other groups (P >0.05). Scale bar: 100 µm. (E) Fluorescence microcopy images of PDOs within NorHA hydrogels at 28 days post-encapsulation, *in vivo* transplantation, and treatment with Verteporfin or DMSO stained for Sox9, Yap, and Ki67. Scale bar: 100 µm. (F, G) Quantification of percentage of (F) Sox9+ cells (%Sox9+) and (G) Ki67+ cells (%Ki67+) per EAC PDO within NorHA hydrogels at 28 days post-encapsulation, *in vivo* transplantation, and treatment with Verteporfin or DMSO (mean ± SEM; (F) n = at least 6 organoids analyzed per group; (G) n = at least 8 organoids analyzed per group). Kruskal-Wallis test with Dunn’s multiple comparisons showed significant differences between 1000 Pa + DMSO and 1000 Pa + Verteporfin, 1000 Pa + DMSO and 100 Pa + DMSO, 1000 Pa + DMSO and 100 Pa + Verteporfin (*P <0.05, ***P <0.001, ****P <0.0001), and no significant differences among other groups (P >0.05). (A-G) Two independent experiments were performed and data are presented for one of the experiments. Every independent experiment was performed with two gels per mouse and five mice per experimental group.

## Discussion

In this study we have established a novel tumor ECM-mimetic hydrogel platform with tunable mechanical properties, controlled presentation of cell-adhesive ligands, and protease-dependent degradation that supports the culture of EAC PDOs. Both mechanical and biochemical properties of the engineered hydrogel were important to enable organoid formation and viability, and we have identified an optimal formulation that supports EAC PDO survival, growth and expansion. Presentation of the cell-adhesive ligands and hydrogel susceptibility to protease-dependent degradation was essential for cell viability and organoid formation, consistent with previous work^38^. In addition, we demonstrated that this engineered hydrogel formulation supports the development of other PDOs, such as Barrett’s esophagus (BE) organoids. This is important in that BE is a major precursor to EAC, thereby permitting elucidation of how matrix mechanics influence progression from a precancer state to a cancer state. Moreover, hydrogel mechanics had direct control over EAC PDO’s fate, eliciting a stem-like behavior via matrix stiffness-mediated activation of the YAP-SOX9 axis. Specifically, we showed that EAC PDOs cultured in NorHA hydrogels with disease-relevant mechanical properties (G’ = 10^3^ Pa) had significant increases in organoid formation and growth, and activation of tumor-associated pathways, as compared to organoids within softer hydrogels. Our observations provide new insights into how cancer cells modulate activity of transcription factors that promote cell proliferation and survival in response to increased ECM stiffness via mechanotransduction pathways. The well-defined engineered hydrogel addresses major limitations of Matrigel™ associated with lot-to-lot variability and inability to uncouple matrix physicochemical properties, establishing the engineered hydrogel as a 3D platform to dissect the pro-tumorigenic contributions of matrix stiffness to EAC development in *in vitro* and *in vivo* settings, thereby addressing a major gap in the field.

Although increased ECM stiffness drives malignant cell transformation, progression and metastasis, current traditional PDX models do not account for tumor ECM mechanics. Therefore, we also exploited the native biocompatibility of the NorHA hydrogel and the susceptibility of EAC PDOs to anti-cancer drugs to establish a novel *in vivo* model for targeted therapy studies. Whilst recent work has focused on understanding the role of engineered-ECM properties in tumor PDO (not EAC) resistance to therapy^23,27–29^, we showed that our novel *in vivo* xenograft model allows for identification of matrix stiffness-dependent expression of transcription factors that can be exploited for targeted therapy in EAC. Therefore, this novel engineered organoid culture platform lays the foundation for application of therapeutics to disrupt contribution of ECM stiffness in EAC, and potentially, other cancers.

## Materials and methods

### Patient sample procurement

Normal and EAC tissue samples from patient biopsies were obtained as paraffin-embedded tissue samples from the Molecular Pathology Shared Resource of the Herbert Irving Comprehensive Cancer Center (HICCC), as approved by Columbia University Institutional Review Board (IRB# AAAS4603; PI: Dr. Julian Abrams). All methods were performed in accordance with Columbia University IRB committee’s regulations on human subject research. All procedures were performed at the New York Presbyterian Hospital/Columbia University Irving Medical Center

### Patient-derived 3D organoid generation and culture

We used a EAC PDO line, namely EAC000, as described previously^13,14^, and a BE PDO line as described previously^41^. Briefly, the human EAC or BE tissue biopsy was digested with Dispase (Corning) and Trypsin-EDTA (Invitrogen), followed by mechanical dissociation and passing through a 100µm cell strainer (Falcon). The enzymes were inactivated by soybean trypsin inhibitor (Sigma-Aldrich), cells were washed in DPBS, counted, and seeded at 25,000 cells per 50µL Matrigel™ (Corning) per one well of a 24-well plate. For passaging of EAC PDOs in Matrigel™, the organoids are dissociated as single-cell suspension with Trypsin–EDTA and seeded at 25,000 cells per 50µL Matrigel™. For passaging of BE PDOs in Matrigel™, the organoids are mechanically dislodged by pipetting through a P200 pipette tip attached to a P1000 pipette tip to break down into small fragments, and seeded at 50-100 fragments per 50µL Matrigel™. As previously described^13^, the organoid growth medium was composed of (50%) L-WRN cell-conditioned medium expressing Wnt-3A, R-Spondin1 and Noggin (WRN), and (50%) Advanced DMEM-F12 (Thermo), supplemented with (1X) GlutaMAX, (10 mM) HEPES, (1X) N-2, (1X) B-27, (1 mM; NAC, (0.5µM) CHIR99021; (250 ng/mL) EGF, (0.5µM) A83-01, (1µM) SB202190, (0.1µM) Gastrin, (20mM) Nicotinamide, (10µM) Y-27632, (10µM) Gentamicin, (1X) Antibiotic-Antimycotic, in (50%). The organoid medium is refreshed every 2-3 days, and organoids are passages every 7 – 14 days. For all experiments, mycoplasma-free PDO lines were limited to fewer than 15 passages.

### Hydrogel synthesis, fabrication, and PDO encapsulation

Hydrogels were synthesized as previously described^36^. Briefly, prior to NorHA macromer synthesis, sodium hyaluronic acid (NaHA; Lifecore Biomedical) was converted to its tetrabutylammonium salt (HA-TBA) using the Dowex 50W proton exchange resin (Millipore-Sigma). To synthesize the NorHA macromer, HA-TBA was dissolved in anhydrous DMSO (2 wt%) with a 3:1 M ratio of 5-norbornene-2-carboxylic acid (mixture of endo and exo isomers; Millipore-Sigma) to HA-TBA repeat units, and 4-(dimethylamino) pyridine (1.5 M ratio to HA-TBA repeat units; Millipore-Sigma) was added under a N2 atmosphere. The product was analyzed by 1H NMR spectroscopy and the NorHA was found to have ∼25% of its repeat units functionalized with norbornene. For fabrication of NorHA hydrogel, NorHA macromer (MW: 30 kDa) was dissolved in DPBS at 2% w/v. Adhesive and crosslinking peptides were custom synthesized by GenScript. Adhesive peptides RGD (GCGYGRGDSPG) or RDG (GCGYGRDGSPG) was dissolved in DPBS at 50 mM (25X final ligand density). Bis-cysteine crosslinking peptide VPM (GCNSVPMSMRGGSNCG) or non-degradable crosslinking agent DTT (1,4-dithiothreitol; Sigma, 3483-12-3) was dissolved in diH2O at 27.3 mM. Photo-initiator, Irgacure 2959 (Ciba, I2959), was dissolved in DPBS at 0.5 wt%. For PDO encapsulation, organoids that were expanded in Matrigel™ for up to 14 days were retrieved by enzymatic digestion of the Matrigel™ using Dispase (Corning) and resuspended at 6.67X final density (final density: 25,000 EAC PDO cells or 50 BE fragments per 25µL hydrogel) in organoid growth medium. Hydrogel precursor solutions and PDO cell solution were mixed and photopolymerized with a curing lamp (OmniCure S1500, Excelitas Technologies) with an internal visible light filter (390 nm) at an intensity of 10 mW/cm^2^ for 5 min.

For all *in vitro* experiments, 25,000 (single-cell suspension) EAC PDO cells or 50 BE fragments were encapsulated in 25 µL NorHA hydrogels. Sample size was established as at least four NorHA hydrogels per condition with the premise that an outcome present in four different hydrogels under a specific condition will reveal the population behavior submitted to this given condition. For all *in vivo* experiments, 200,000 cells (single-cell suspension of EAC PDOs) were encapsulated in 50 µL NorHA hydrogels. To collect organoids from hydrogels for downstream assays, we transfer the NorHA hydrogels to 1mg/ml hyaluronidase (Sigma, H3884), or Matrigel™ to Dispase (Corning), and incubate at 37°C for 20 minutes to digest the hydrogel and release the organoids.

### Rheological characterization

Storage moduli (G’) were characterized using an oscillatory shear rheometer (AR2000, TA Instruments) fitted with a 20 mm diameter cone and plate geometry and 27 μm gap. Time sweeps (0.5% strain, 1 Hz) were performed at 37°C to characterize bulk gelation upon exposure to visible light filter (390 nm) at an intensity of 10 mW/cm^2^ for 10 min using OmniCure S1500 lamp (Excelitas Technologies).

### Viability assay and quantification

NorHA hydrogels were incubated in 2 µM Calcein-AM (Life Technologies), in growth medium for 1 h. Samples were imaged using Celigo Image Cytometer (Nexcelom). Quantification of viability was performed by calculating the percentage of PDOs that stained positive for Calcein-AM using ImageJ (National Institute of Health, USA). The results are representative of three independent experiments performed with four NorHA hydrogel or Matrigel™ samples per experimental group.

### Quantitative Reverse Transcription PCR

RNA isolation was achieved using the RNAqueous™ Phenol-free total RNA Isolation kit (Invitrogen, AM1912), according to manufacturer’s instructions. cDNA was synthesized using the Applied Biosystems™ High-Capacity cDNA Reverse Transcription Kit (Fisher Scientific, 43-688-13) according to manufacturer’s instructions. Quantitative PCR was performed using the Applied Biosystems 7500 Real Time PCR System. The primer sequences used were: SOX9 forward sequence ACTTGCACAACGCCGAG and SOX9 reverse sequence CTGGTACTTGTAATCCGGGTG; YAP forward sequence AATTGAGAACAATGACGA and YAP reverse sequence AGTATCACCTGTATCCATCTC; P53 forward sequence CTTCCATTTGCTTTGTCCCG and P53 reverse sequence CATCTCCCAAACATCCCTCAC; STAT3 forward sequence GGTACATCATGGGCTTTATC, and STAT3 reverse sequence TTTGCTGCTTTCACTGAATC; housekeeping gene, YWHAZ forward sequence ACTTTTGGTACATTGTGGCTTCAA; housekeeping gene, YWHAZ reverse sequence CCGCCAGGACAAACCAGTAT. Gene expression of all samples was normalized to its corresponding housekeeping gene expression before normalization to control sample.

### Immunofluorescence and immunohistochemistry analysis

For immunofluorescence or immunohistochemistry staining of paraffin sections from PDOs, xenograft implants or human tissue samples, these were fixed with 4% (w/v) paraformaldehyde at 4°C for 4 hr to overnight. Sections were deparaffinized and heat-induced antigen retrieval was performed using 10 mM citric acid buffer (pH 6) for 15 min. Samples were permeabilized using 0.5% (w/v) Triton X-100 for 10 min and blocked with 5% donkey serum. Primary antibody incubation was performed overnight at 4°C at a 1:100 dilution, unless stated otherwise below. Secondary antibody incubation was performed for 30 min at 37°C at a 1:200 dilution. The following primary antibodies were used: CD44 (1:200 dilution; Cell Signaling, 3570S), FN1 (Cell Signaling, 26836S), Hyaluronic Acid Binding Protein (HABP; Millipore-Sigma, 385911), Ki67 (1:50 dilution; BD Biosciences, 550609), YAP1 (Cell Signaling, 14074), Sox9 (1:20 dilution; R&D Systems, AF3075), MUC5ac (1:200 dilution; Cell Signaling, 61193). DAPI (Vector Laboratories, H-1500) was used as counterstain for immunofluorescence, and Hematoxylin stain (Leica, 3801560) was used as counterstain for immunohistochemistry. The following secondary antibodies were used: Alexa Fluor 488 donkey anti-goat (A11055), Alexa Fluor 488 donkey anti-mouse IgG (Invitrogen, A32766), Alexa Fluor 555 donkey anti-rabbit (A32794).

### Image acquisition and quantification

Brightfield images of PDOs were acquired using the Celigo Image Cytometer. Quantification of organoid size and density was be performed using Celigo Image Cytometer and its analytical algorithms. Acquisition of fluorescence, immunohistochemistry, and H&E images was performed using Keyence BZ-X800 Cell Imaging Microscope. Quantification of percentage of fluorescently labeled cells (e.g. nuclear YAP+ cells, Fig. 4e) was done using Keyence BZ-X800 Cell Imaging Microscope and its analytical algorithms.

### Animal models

All animal studies were conducted following approved protocols established by Columbia University’s Institutional Animal Care and Use Committee (IACUC) in accordance with the US Department of Agriculture (USDA), Animal and Plant Health Inspection Service (APHIS) regulations, and the National Institutes of Health (NIH) Office of Laboratory Animal Welfare (OLAW) regulations governing the use of vertebrate animals. Male (8 weeks old) NOD-scid IL2Rg-null (NSG) mice (Jackson Laboratory) were used for all experiments.

### Xenograft transplantation

Single-cell suspension of EAC PDOs were embedded in NorHA hydrogels two hours prior to subcutaneous transplantation in the back (flanks) of male NOD-scid IL2Rg-null (NSG) mice (Jackson Laboratory). Mice were anesthetized by intraperitoneal (IP) injection of ketamine (100 mg/kg)/xylazine (10 mg/kg) solution and hair was removed from the back to expose skin. A small incision was made through the skin on the mouse’s right and left flanks, and the connective tissue cleared to make a small subcutaneous space (pocket) on each side. One NorHA hydrogel containing EAC cells was delivered to each subcutaneous pocket using a surgical spatula (1 implantation per pocket, 2 pockets per animal). The skin was closed using absorbable sutures. The mice were euthanized and the transplant retrieved after 4 weeks. For drug treatment experiments, 1 week after transplantation, Verteporfin (100mg/kg in DMSO; maximum of 150 µL per injection) or DMSO treatment (150 µL per injection) was given every other day for 3 weeks. At the end of 3 weeks of treatment, mice were euthanized and transplant retrieved. The results are representative of two experiment performed with five mice per condition (one organoid implanted per subcutaneous pocket). Sample size was established as 10 transplants (2 per mouse) with the premise that an outcome present in 10 different samples under a specific condition will reveal the population behavior submitted to this given condition. No statistical method was used to predetermine sample size.

### Tumor vs normal RNA-sequencing analysis

Analysis of RNA-sequencing (RNA-seq) data was performed using a previously published online tool^35^ collecting data from The Cancer Genome Atlas (TCGA) and The Genotype Tissue Expression Project (GTEx). The esophageal carcinoma (“ESCA”) dataset was used. The data were plotted on a log scale (log2(TPM + 1)) with a jitter size of 0.4. “Match TCGA normal and GTEx data” was selected.

### Statistical analysis and reproducibility

All experiments were performed three or more times independently under similar conditions, except experiments shown in Figs. 4 and 5, which were performed twice. Plots shown are of one experiment representative of all independent experiments performed under similar conditions. All immunofluorescence or immunohistochemistry images shown are representative of at least 20 images that were stained and imaged for each specific marker per experimental group for each independent experiment. All statistical analyses were performed using GraphPad Prism 6.0. For statistical comparisons between two groups a Welch’s t-test (or Mann-Whitney test for nonparametric data) was used, and among more than two groups a one-way ANOVA (or Kruskal-Wallis test for nonparametric data) was used. For all data, P <0.05 was considered statistically significant. P values of statistical significance are represented as ****P <0.0001, ***P<0.001, **P<0.01, *P<0.05. To ensure rigor and reproducibility, other colleagues not involved in this study masked the labels of prepared tissue slides (histology/immunofluorescence) prior to image acquisition. For all experiments, mycoplasma-free PDO lines were limited to fewer than 15 passages.

## Acknowledgements

This research was supported by the National Institute of Health (A.K.R. acknowledges NCI P01-CA098101 and NCI U54-CA163004) and the National Science Foundation through the Center for Engineering Mechano-Biology STC (J.A.B. acknowledges CMMI: 15-48571). R.C.-A. acknowledges the Charles H. Revson Senior Fellowship in Biomedical Science. We thank the Molecular Pathology Shared Resource of the HICCC, and Drs. Hanina Hibshoosh and Julian A. Abrams for their assistance in procuring patient tissue samples.

## Author contributions

R.C.-A. conducted all experiments, collected data and performed data analyses. S.W.K. assisted with data collection and experimental analyses. K.S. assisted with *in vivo* experiments. C.L. assisted with *in vitro* experiments and rheological measurements. J.T.G. assisted with data collection. R.C.-A., T.K., J.A.B. and A.K.R. conceptualized and designed the project and experiments. R.C.-A., J.A.B. and A.K.R. wrote the manuscript.

## Data availability

All data needed to evaluate the conclusions in the paper are present in the manuscript and/or the Supplementary Information. Raw data related to this paper are available for research purposes upon reasonable request.

