## Supplementary Figures for "Engineered hydrogel reveals contribution of matrix mechanics to esophageal adenocarcinoma 3D organoids and identify matrix-activated therapeutic targets"

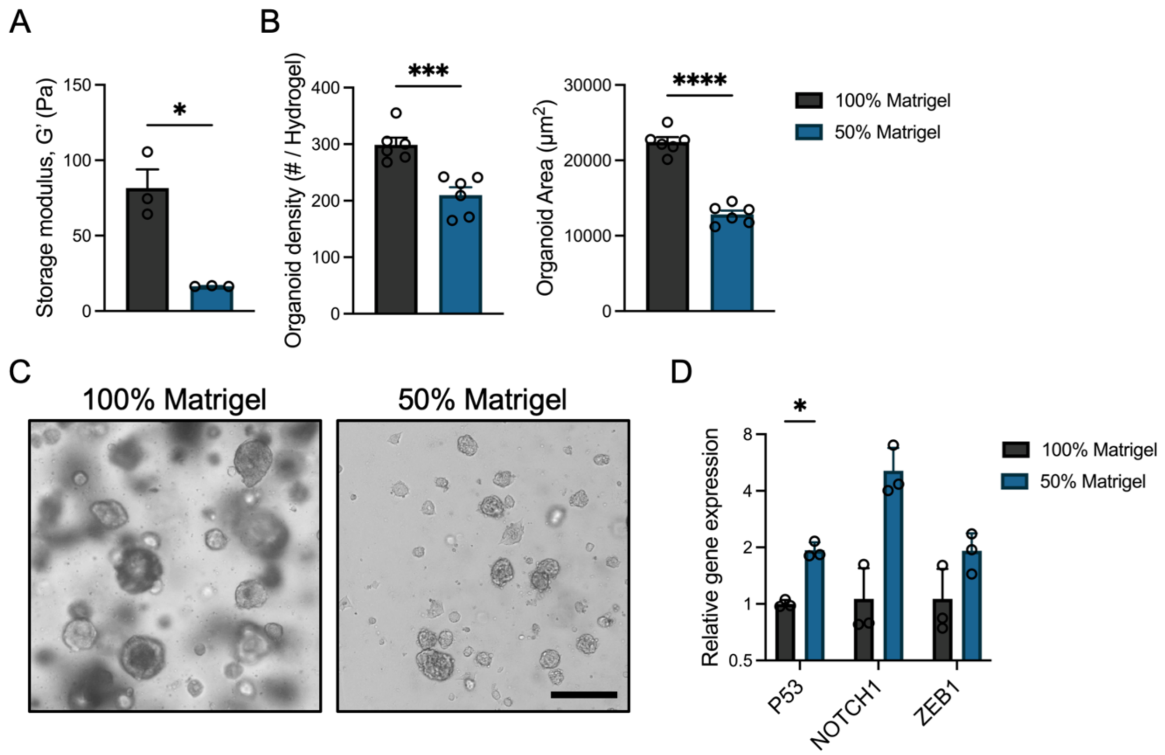


**Supplementary Figure 1: EAC PDOs growth and gene expression in different Matrigel™ concentrations.** (A) Relationship between Matrigel™ concentration (%) and storage modulus, G' (mean ± SEM; n = 3 independently prepared hydrogels per condition). Welch's t-test with two-tailed comparison showed significant differences between 100% and 50% Matrigel™ (*P <0.05). (B) Quantification of PDO density and size (area) as a function of Matrigel™ concentration at 10 days post-encapsulation (mean ± SEM; n = 6 hydrogels per group). Welch's t-test with two-tailed comparison showed significant differences between 100% and 50% Matrigel™ (***P <0.001, ****P <0.0001). (C) Representative transmitted light images of EAC PDOs in 100% or 50% Matrigel™. Scale bar: 250 µm. (D) Relative gene expression levels of EAC PDOs in 100% or 50% Matrigel™. RNA levels of EAC-associated genes (P53, NOTCH1, ZEB1), as quantified by RT-qPCR (mean ± SEM; n = 3 samples per group), and normalized to 100% Matrigel™. Multiple Welch's t-test were used to identify statistical differences (*P <0.05). (A-D) Three independent experiments were performed and data are presented for one of the experiments. Every independent experiment was performed with six gel samples per experimental group.

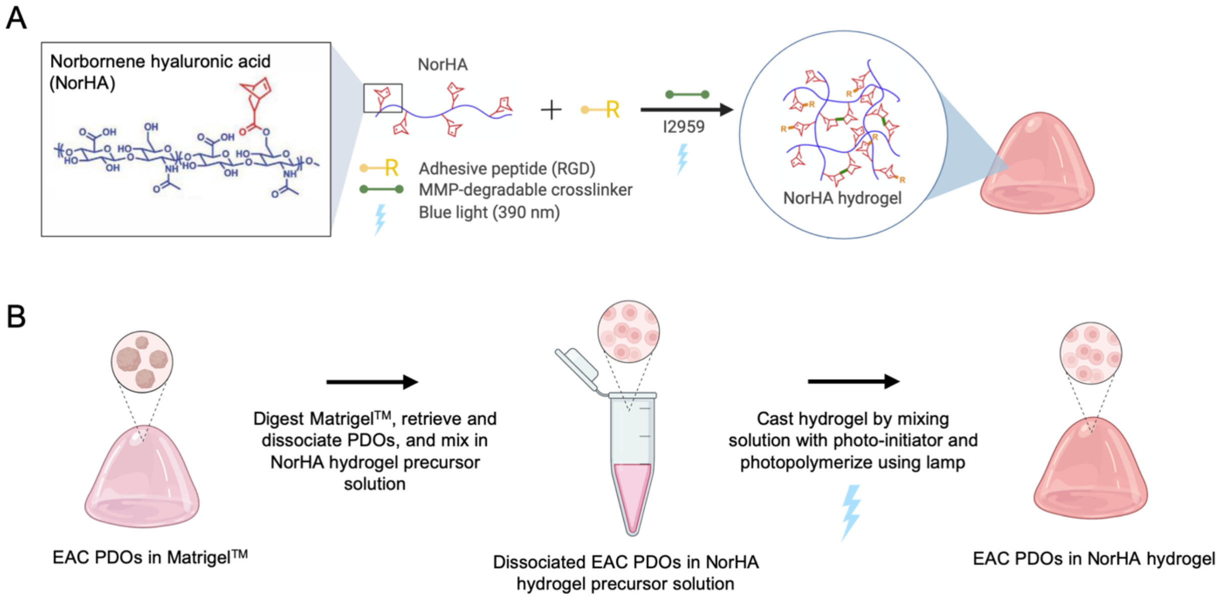


**Supplementary Figure 2: NorHA hydrogel fabrication.** (A) Schematic of NorHA hydrogel fabrication process to cast gels through the light-initiated thiol-ene reaction among a mono-thiol RGD adhesive ligand, a di-thiol MMP-degradable crosslinker and NorHA. Adapted from Gramlich WM, Kim IL, Burdick JA. Synthesis and orthogonal photopatterning of hyaluronic acid hydrogels with thiol-norbornene chemistry, Biomaterials (2013). (B) Schematic of EAC PDO encapsulation within NorHA hydrogel. Created with BioRender.com.

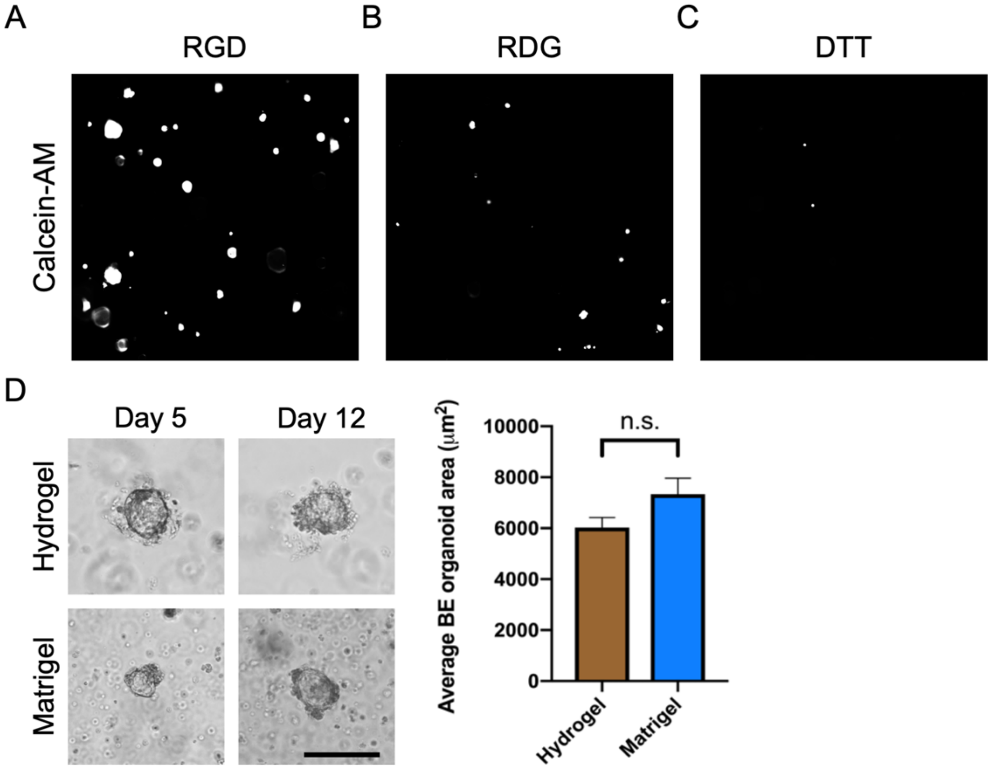


**Supplementary Figure 3: Engineered NorHA hydrogel presenting RGD adhesive peptide and MMP-degradable crosslinker promote EAC PDO viability and BE PDO growth.** (A-C) Representative fluorescence images of EAC PDO viability (Calcein-AM) at 7 days after encapsulation within NorHA (100 Pa) hydrogels functionalized with (A) RGD, or (B) RDG, or (C) functionalized with RGD and crosslinked with non-degradable agent DTT. Scale bar, 500 μm. (D) Representative transmitted light images and quantification of size (area) of BE PDOs cultured in engineered NorHA hydrogel or Matrigel™ at day 12 post-encapsulation (mean ± SEM; n = at least 50 organoids analyzed across 4 hydrogels per group). Welch's t-test with two-tailed comparison showed no significant differences between groups (ns = P > 0.05). Scale bar: 200 μm. (A-D) Three independent experiments were performed and data are presented for one of the experiments. Every independent experiment was performed with four gel samples per experimental group.

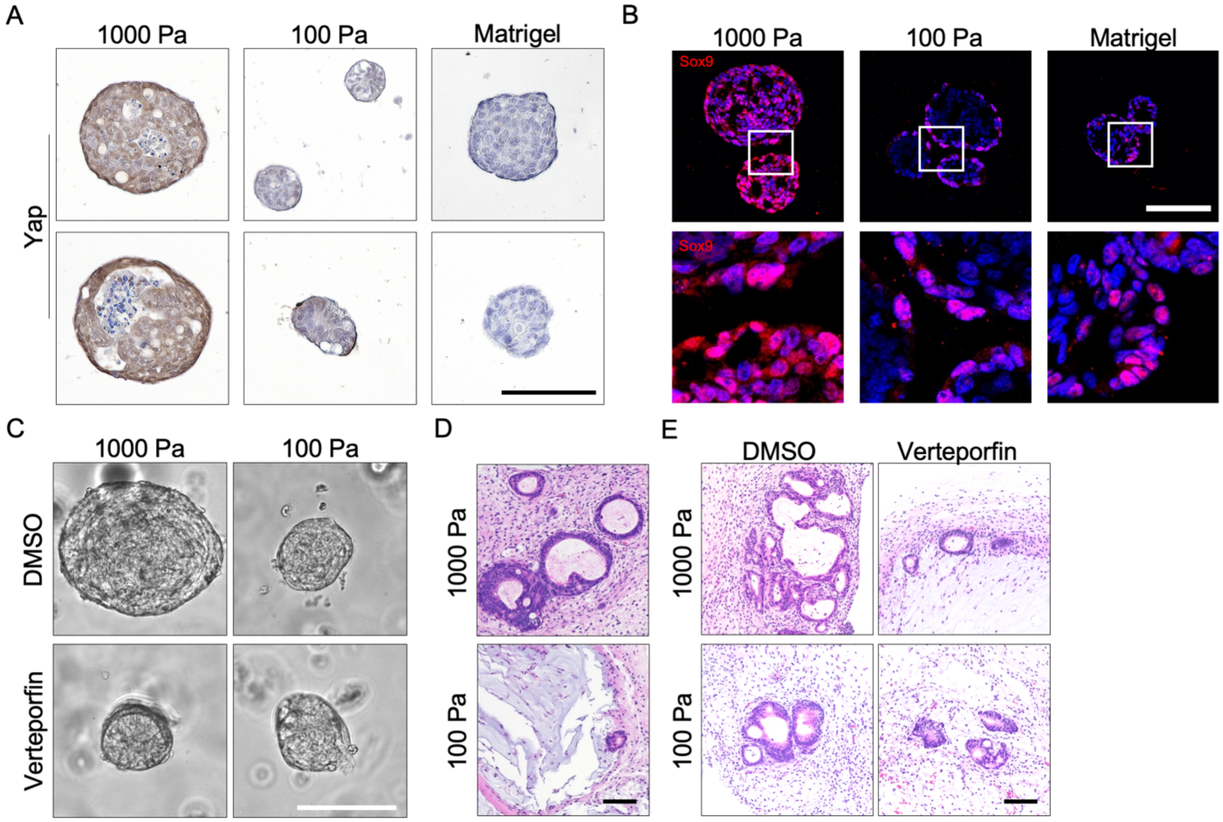


**Supplementary Figure 4: Matrix stiffness-mediated growth and activation of the Yap-Sox9 axis in EAC PDOs in *in vitro* and *in vivo* models.** (A) Representative images of (A) immunohistochemical staining of Yap, or (B) immunofluorescence staining of Sox9 to EAC PDOs cultured within NorHA hydrogels of different stiffnesses at 14 days post-encapsulation. (C) Representative transmitted light images of EAC PDOs cultured within NorHA hydrogels of different stiffnesses and treated with 5 nM Verteporfin or DMSO. (D) Representative histological (H&E) images of EAC PDOs cultured within NorHA hydrogels at 28 days post-encapsulation and *in vivo* transplantation. (E) Representative histological (H&E) images of EAC PDOs cultured within NorHA hydrogels at 28 days post-encapsulation and *in vivo* transplantation, and treated with 5 nM Verteporfin or DMSO. Scale bars: 100 μm. (A-C) Three independent experiments were performed and data are presented for one of the experiments. Every independent experiment was performed with four gel samples per experimental group. (D,E) Two independent experiments were performed and data are presented for one of the experiments. Every independent experiment was performed with two gels per mouse and five mice per experimental group.

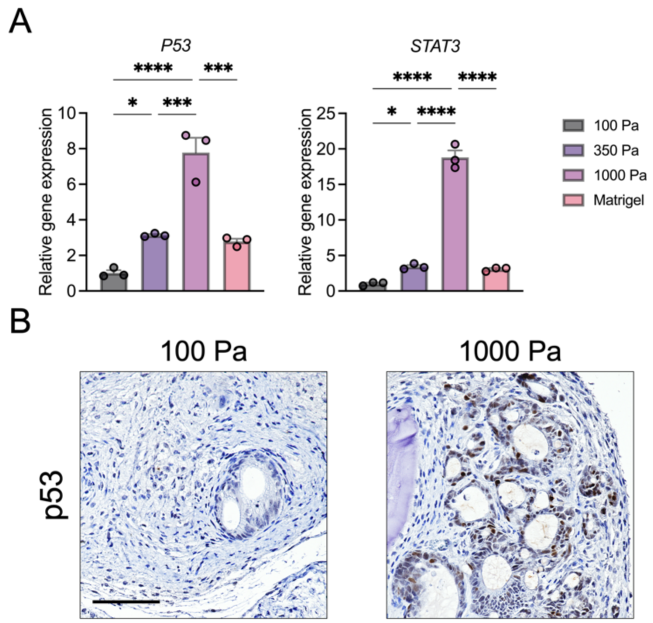


**Supplementary Figure 5: Matrix stiffness-mediated expression of EAC-associated genes.** (A) Transcriptional expression of P53 and STAT3 in EAC PDOs within NorHA hydrogels of different stiffness at 14 days post-encapsulation (mean ± SEM; n = 3 technical replicates; RNA levels normalized to 100 Pa; representative of 1 independent experiment). One-way ANOVA with Tukey's multiple comparisons test showed significant differences between 100 Pa and 350 Pa, 100 Pa and 1000 Pa, 350 Pa and 1000 Pa, 1000 Pa and Matrigel$^{TM}$ (*P <0.05, ***P <0.001, ****P <0.0001). Three independent experiments were performed and data are presented for one of the experiments. Every independent experiment was performed with four gel samples per experimental group. (B) Representative images of immunohistochemical p53 staining of EAC PDOs cultured within NorHA hydrogels of different stiffnesses at 28 days post-encapsulation and *in vivo* transplantation. Scale bar: 100 µm. Two independent experiments were performed and data are presented for one of the experiments. Every independent experiment was performed with two gels per mouse and five mice per experimental group.
